# A non-canonical metal center drives activity of the *Sediminispirochaeta smaragdinae* metallo-β- lactamase SPS-1

**DOI:** 10.1101/336024

**Authors:** Zishuo Cheng, Jamie VanPelt, Alexander Bergstrom, Christopher Bethel, Andrew Katko, Callie Miller, Kelly Mason, Erin Cumming, Huan Zhang, Robert Kimble, Sarah Fullington, Stacey Lowery Bretz, Jay C. Nix, Robert A. Bonomo, David L. Tierney, Richard C. Page, Michael W Crowder

**Affiliations:** Department of Chemistry and Biochemistry, 651 E. High Street, 160 Hughes Laboratories, Miami University, Oxford, OH 45056; Research Service, Louis Stokes Cleveland Department of Veterans Affairs Medical Center, Cleveland, OH 44106; Departments of Medicine, Pharmacology, Molecular Biology and Microbiology, Biochemistry, Proteomics and Bioinformatics, and the CWRU-Cleveland VAMC Center of Antimicrobial Resistance and Epidemiology, Cleveland, OH 44106; Molecular Biology Consortium, Beamline 4.2.2, Advanced Light Source, Lawrence Berkeley National Laboratory, Berkeley, CA 94720

## Abstract

In an effort to evaluate whether a recently reported putative metallo-β-lactamase (MβL) contains a novel MβL active site, SPS-1 from *Sediminispirochaeta smaragdinae* was over-expressed, purified, and characterized using spectroscopic and crystallographic studies. Metal analyses demonstrate that recombinant SPS-1 binds nearly 2 equivalents of Zn (II), and steady-state kinetic studies show that the enzyme hydrolyzes carbapenems and certain cephalosporins but not β-lactam substrates with bulky substituents in the 6-7 position. Spectroscopic studies on Co (II)-substituted SPS-1 suggest a novel metal center in SPS-1, with reduced spin coupling between the metal ions and a novel Zn_1_ metal binding site. This site was confirmed with a crystal structure of the enzyme. The structure shows a Zn_2_ site that is similar that that in NDM-1 and other subclass B1 MβLs; however, the Zn_1_ metal ion is coordinated by 2 histidine residues and a water molecule, which is held in position by a hydrogen bond network. The Zn_1_ metal is displaced nearly 1 Å from the position reported in other MβLs. The structure also shows extended helices above the active site, which create a binding pocket that precludes the binding of substrates with large, bulky substituents in the 6/7 position of β-lactam antibiotics. This study reveals a novel metal binding site in MβLs, and suggests that the targeting of metal binding sites in MβLs with inhibitors is now more challenging with the identification of this new MβL.

## INTRODUCTION

*β*-Lactam antibiotics, such as penicillins, cephalosporins, and carbapenems, have long served as treatment for bacterial infections by inactivation of bacterial transpeptidases. These compounds constitute the largest class of clinically-available antibiotics. In response to wide use (and overuse) of β-lactam containing drugs, bacteria have evolved to produce β-lactamases, which hydrolyze the β-lactam bond and render the antibiotics inactive.

Of the four classes of β-lactamases (A, B, C, & D), only class B enzymes require 1-2 Zn (II) ions (rather than an active site Ser) for catalysis and are termed metallo-*β*-lactamases (M*β*Ls) (1-3). M*β*Ls adopt an α*ββ*α fold, and among the three subclasses (B1, B2, & B3), B1 and B3 MβLs generally contain a dinuclear Zn (II) active site, while B2 M*β*Ls are active with mononuclear Zn (II) sites. The B1 M*β*Ls coordinate one Zn (II) (in the Zn_1_ site) with three histidine residues, while the Zn (II) in the Zn_2_ site is bound to one cysteine, one aspartate, one histidine, and a terminally-bound solvent molecule. The Zn (II) ions are bridged by a solvent molecule, resulting in a Zn—Zn distance of 3.3-3.9 Å (2,4). The B3 MβLs possess the same Zn_1_ site as the B1 MβLs; however, in the Zn_2_ site, the cysteine ligand is replaced with an additional His residue. Subclass B1 contains the largest number of clinically-relevant MβLs, including VIMs (Verona integrin-encoded MβL), IMPs (imipenemase), and NDMs (New Delhi MβL). Most recent studies have focused on MβLs found in pathogenic strains, however the risk of transfer of antibiotic resistance determinants from a non-pathogenic strain to a more clinically-relevant organism necessitates the need to study MβLs not yet found in known human pathogens (5).

Recently, with the use of the Markov model and a bioinformatic study, Berglund *et al.* identified 279 unique, potential subclass B1 MβL genes, and 76 of the corresponding gene products were novel and able to hydrolyze imipenem (6). One gene, *SPS-1*, was found in the bacterium *Sediminispirochata smaragdinae*, (NCBI Reference sequence WP_013255389.1). The *S. smaragdinae* strain SEBR 4228^T^ was discovered in a water sample near a Congo offshore oilfield (7). SEBR 4228^T^ is Gram-negative, chemoorganotrophic and anaerobic bacterium that adopts a spiral, corkscrew-like structure. This organism is halophilic and able to grow in high-salt environments. SPS-1 was predicted to be a MβL; however, SPS-1 lacked the typical H-X-H-Y-D motif found in all MβLs except the B2 MβLs. In SPS-1, the first consensus histidine residue was substituted with a glycine (6). In an effort to determine if SPS-1 is indeed a MβL and how the histidine to glycine substitution affected the consensus Zn_1_ site, we cloned, over-expressed, and purified the enzyme, and characterized the recombinant protein with kinetic, spectroscopic, and crystallographic studies.

## RESULTS

**Phylogenetic analysis of SPS-1.** MβLs can be grouped into three subclasses phylogenetically (1). Subclass B1 MβLs contain a Zn_1_ binding site with three histidine residues (His116, His118, His196) and a Zn_2_ site with three other residues (Asp120, Cys221, His263) (1-3). Subclass B2 MβLs contain a Zn_1_ site with one altered residue (Asn116, His118, His196) and Zn_2_ site identical to that of the subclass B1 MβLs; the altered Zn_1_ site results in B2 MβLs exhibiting activity as mononuclear Zn (II) enzymes (8,9). The subclass B3 MβLs have a varied Zn_1_ binding site (His/Gln116, His118, His196) and a distinctive Zn_2_ binding site, which does not contain a cysteine residue (Asp120, His121, His263). To determine the phylogenetic position of SPS-1 in the classification of MβLs, phylogenetic analysis was conducted using Phylogeny (10). The analyses showed that enzymes in subclasses B1 and B2 descended from a common ancestor and that SPS-1 was grouped in the B1 class and is highly similar to the MβL SPM-1 (Figure 1) (11). Subgroup B3 MβLs are divergent from subgroups B1 and B2.

**FIGURE 1.**
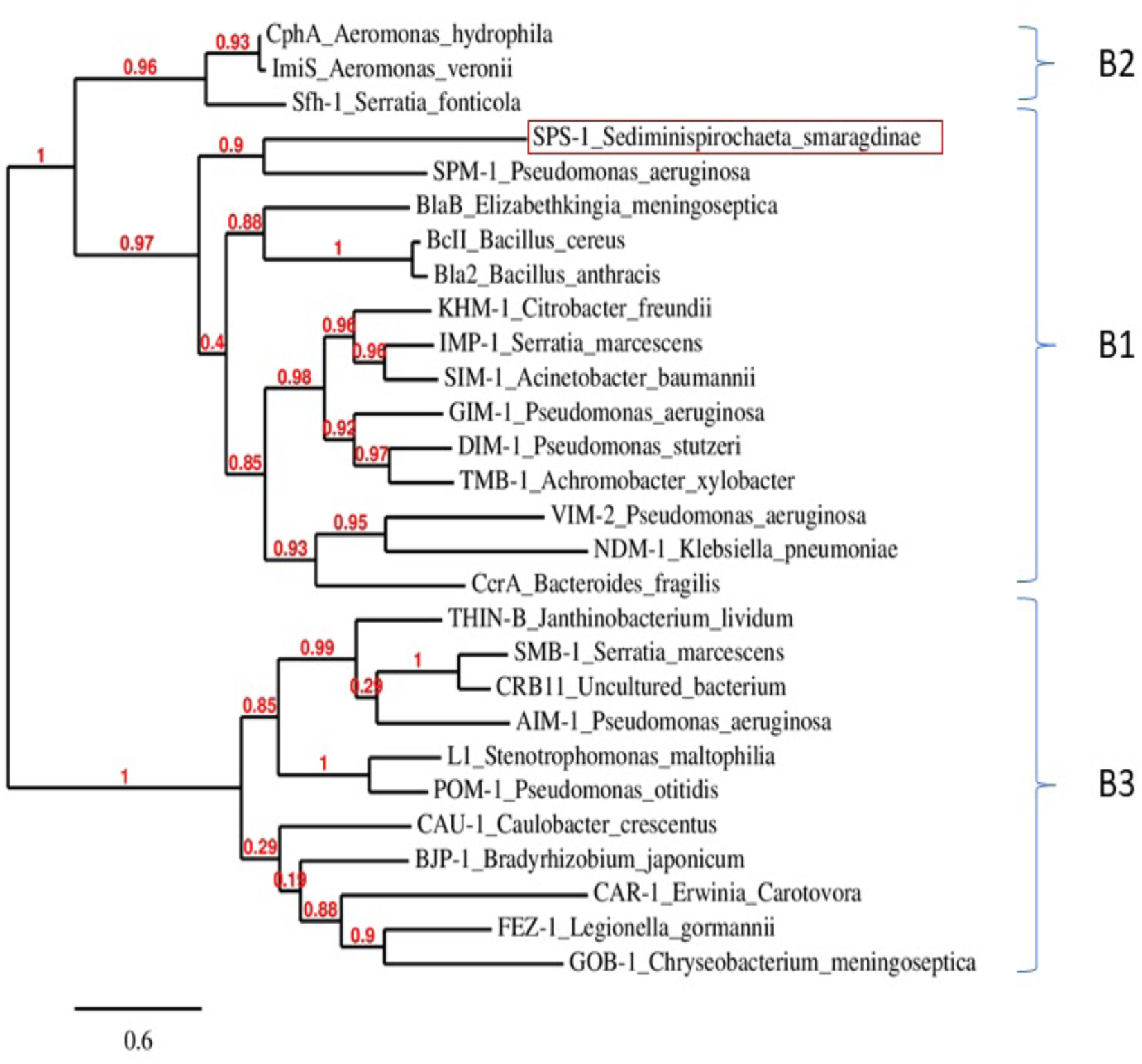
The phylogenetic tree of class B metallo-β-lactamase sequences. The protein sequences of SPS-1 and other selective metallo-β-lactamase were download and analyzed by Phylogeny.fr. The sequence include SPS-1 (WP_013255389.1), BcII (M11189), CcrA (M63556), IMP-1 (S71932), VIM-2 (AF191564), BlaB (AF189298), SPM-1 (AY341249), NDM-1 (JN420336), GIM-1 (AJ620678), SIM-1 (AY887066), DIM-1 (GU323019), TMB-1 (FR771847), Bla2 (CP001598), KHM-1 (AB364006), CphA (X57102), Sfh-1 (AF197943), ImiS (Y01415), L1 (AB294542), FEZ-1 (Y17896), BJP-1 (NP772870), AIM-1 (AM998375), THIN-B (CAC33832), GOB-1 (ABO21417), CAU-1 (CAC87665), CAR-1 (Q6D395), SMB-1 (AB636283), POM-1 (ADC79555), CRB11 (ACS83724). The line segment with the number ‘0.6’ shows the length of branch that represents an amount genetic change of 0.6.

**Over-expression, purification, and characterization of ZnZn-SPS-1.** A region of the gene for SPS-1 corresponding to residues 32-276 was synthesized to aid in cloning SPS-1 into a pET28A overexpression vector. The DNA sequence corresponding to the first 31 amino acids was not included as these residues were predicted by Phobius (12) to comprise a signal peptide. Over-expressed SPS-1 was purified with a single-step HisTrap column. TEV protease was used to remove the His_6_-tag, and a second HisTrap column was used to separate SPS-1 from the cleaved tag and TEV protease. The purity of the protein was shown to be >95% by SDS-PAGE. The predicted molecular weight for tag-free SPS-1 is 27466.11 Da, as determined by the ExPASy-Compute pI/Mw tool (13). The MALDI-TOF mass spectrum of tag-free SPS-1 showed a single peak at 27,469 ± 267 m/z (Figure S1), which demonstrated that the protein was purified as a single polypeptide. The yield for tag-free SPS-1 was >20 mg protein per liter of LB media. The tertiary structure of recombinant SPS-1 was evaluated using fluorescence spectroscopy. The resulting spectrum showed a single peak at 345 nm (excitation of 280 nm), suggesting that the majority of the protein consisted of one folded conformation (data not shown) (14). ICP-AES revealed that SPS-1 binds 1.9 ± 0.1 equivalents of Zn (II), similar to the metal content of NDM-1 and other well-characterized dinuclear Zn (II)-containing MβLs (4,15).

**Steady-state kinetics.** Steady-state kinetic studies were conducted at 37 °C using chromacef, ampicillin, penicillin G, imipenem, meropenem, cephalothin, cefuroxime, and cephalexin as substrates. The resulting kinetic data are shown in Table 1. When using chromacef and cephalothin as the substrate, SPS-1 exhibited *K*_m_ values of 120 ± 9 μM and 114 ± 15 μM and *k*_cat_ values of 2.6 ± 0.1 s^-1^ and 3.1 ± 0.1 s^-1^, respectively. When using imipenem and meropenem as substrates, SPS-1 exhibited *K*_m_ values of 288 ± 48 μM and 168 ± 33 μM and *k*_cat_ values of 471 ± 58 s^-1^ and 519 ± 49 s^-1^, respectively. When ampicillin, penicillin G, cefuroxime, and cephalexin were used as substrates, no activity was observed. Compared to NDM-1 and SPM-1, the *K*_m_ values with different substrates exhibited by SPS-1 are much larger, resulting in lower catalytic efficiencies (Table 1). In contrast to most other MβLs (1,2), SPS-1 exhibits substrate specificities for carbapenems and certain cephalosporins.

**Table 1:** Steady-State Kinetic Parameters for different β-lactams by SPS-1, NDM-1 and SPM-1.

| Substrate | SPS-1 |  |  | SPM-1(11) |  |  | NDM-1 (17) (53) |  |  |
| --- | --- | --- | --- | --- | --- | --- | --- | --- | --- |
| | $K_m(\mu\text{M})$ | $k_{\text{cat}}(\text{s}^{-1})$ | $k_{\text{cat}}/K_m$<br>($\text{s}^{-1}/\mu\text{M}$) | $K_m$<br>($\mu\text{M}$) | $k_{\text{cat}}$<br>($\text{s}^{-1}$) | $k_{\text{cat}}/K_m$<br>($\text{s}^{-1}/\mu\text{M}$) | $K_m(\mu\text{M})$ | $k_{\text{cat}}$<br>($\text{s}^{-1}$) | $k_{\text{cat}}/K_m$<br>($\text{s}^{-1}/\mu\text{M}$) |
| chromacef | 120±9 | 2.6±0.1 | 0.021 | Not available | | | 5.1±0.9 | 4.2 | 0.8 |
| ampicillin | Not observed | | | 72±3 | 117±2 | 1.6 | 22 | 15 | 0.66 |
| penicillin G | Not observed |  |  | Not available |  |  | 16 | 11 | 0.68 |
| imipenem | 288±48 | 471±58 | 1.63 | 37±4 | 33±2 | 1 | 94 | 20 | 0.21 |
| meropenem | 168±33 | 519±49 | 3 | 281 ±<br>27 | 63 ± 3 | 0.22 | 49 | 12 | 0.25 |
| cephalothin | 114±15 | 3.1 ±0.1 | 0.027 | 4±1 | 43±2 | 11.7 | 10 | 4 | 0.4 |
| cefuroxime | Not observed | | | 4 ± 1 | 37 ± 3 | 8.8 | 8 | 5 | 0.61 |
| cephalexin | Not observed |  |  | Not available |  |  | Not available |  |  |

**Isothermal titration calorimetry (ITC).** ITC was used to determine if SPS-1 binds to *L*-captopril, which has been reported to be a competitive inhibitor of several MβLs (16). Previous studies have shown that the S1 atom in *L*-captopril inserts directly between the metal ions in NDM-1 (16), displacing the bridging water-hydroxide. The crystal structure of the NDM – 1-captopril complex showed that *L*-captopril has two binding faces: a hydrophobic face that interacts with the Val73 and Met67 of the L3 loop and a hydrophilic face that hydrogen bonds to Asn220 on the L10 loop (16). In our experimental conditions, *L*-captopril binds to NDM-1 with a K_d_ value of 1.95 μM (Figure 2), while there is no binding between SPS-1 and *L*-captopril up to concentrations of 50 μM *L*-captopril. This result suggests a different active site configuration in SPS-1 as compared to NDM-1.

**FIGURE 2.**
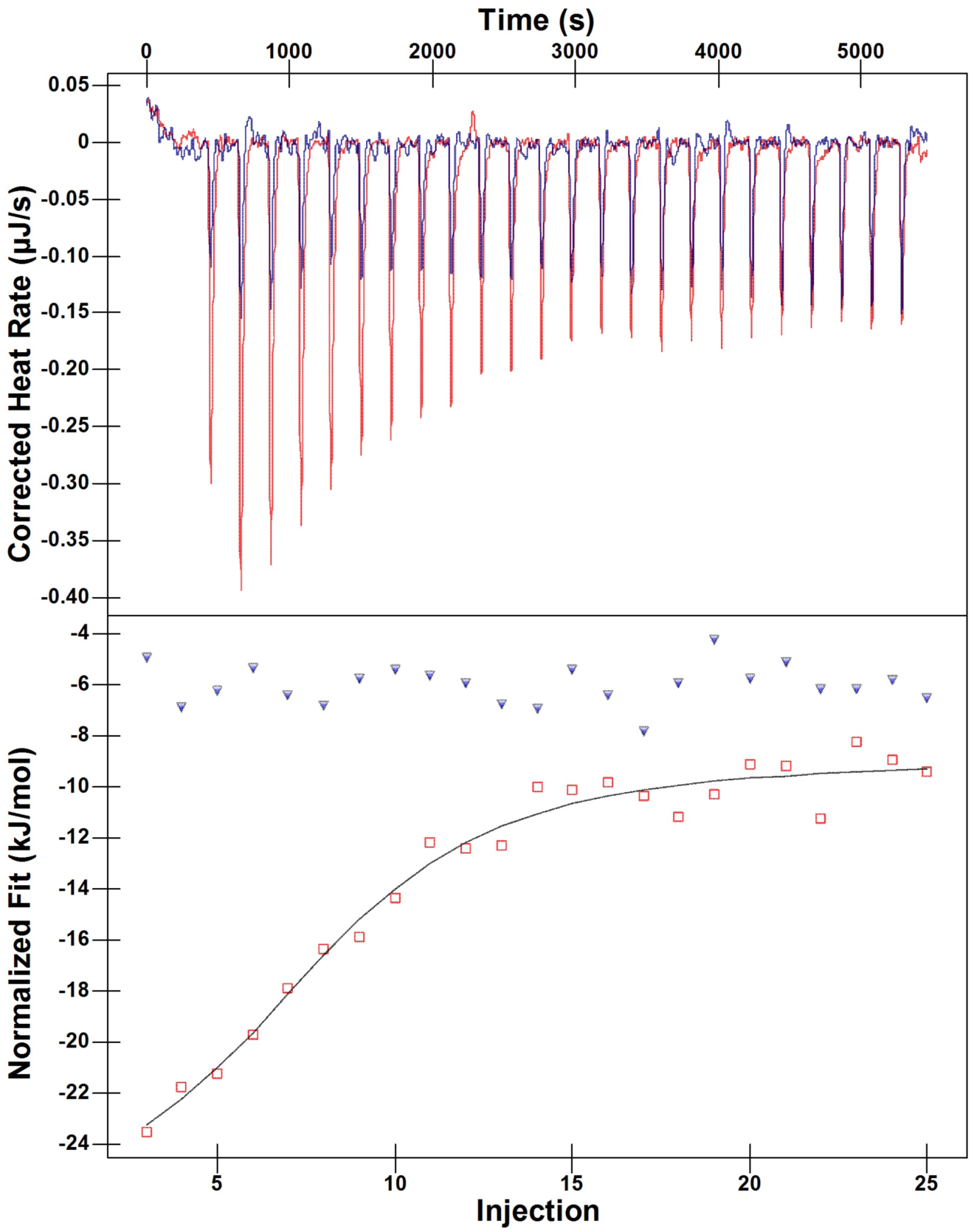
ITC results of *L*-captopril titrations with NDM-1 and SPS-1. Upper panels show the titration thermograms, and lower panels the data integration with fitted curves. NDM-1 protein is highlighted with red, and SPS-1 is highlighted with blue.

**UV-Vis spectroscopy.** To obtain information on the metal binding sites in SPS-1, UV-Vis spectra of Co (II)-substituted SPS-1 were recorded, and Co (II)-substituted NDM-1 was used as a control (17). Metal-free SPS-1 and NDM-1 were prepared as described in Experimental procedures. *Tris* (2-Carboxyethyl)phosphine) (TCEP, 1 mM) was added to metal-free SPS-1, prior to adding Co (II), in order to prevent Co (II) oxidation (17). Glycerol (10%) was added to the metal-free SPS-1 before Co (II) addition to prevent protein precipitation (18). As previously reported (15,17), Co (II)-substituted NDM-1 exhibited an intense peak at 330 nm, which was previously attributed to a Cys-S to Co (II) ligand-to-metal charge transfer transition (19-22). The other peaks at 330, 510, 549, 615, and 640 nm (Figure 3) were assigned to ligand field transitions of high-spin Co (II) exhibiting a Jahn-Teller distortion (19,23) The extinction coefficients of these ligand field transitions are consistent with one Co (II) having a coordination number of 4 and the other having a coordination number of 5 (15). Conversely, the spectra of Co (II)-substituted SPS-1 revealed a sharp absorption at 346 nm (Figure 3), suggesting a similar Zn_2_ site in SPS-1 and NDM-1. However, in the spectra of Co (II)-substituted SPS-1, the ligand field transitions between 500 and 650 nm are broader and lack distinct features. Importantly, the extinction coefficient of this peak is 56 M^-1^cm^-1^ (or about 28 M^-1^cm^-1^ per Co (II)), suggesting that both Co (II) ions are 6-coordinate (15). A similar spectrum was previously reported for Co (II)-substituted L1 (24).

**FIGURE 3.**
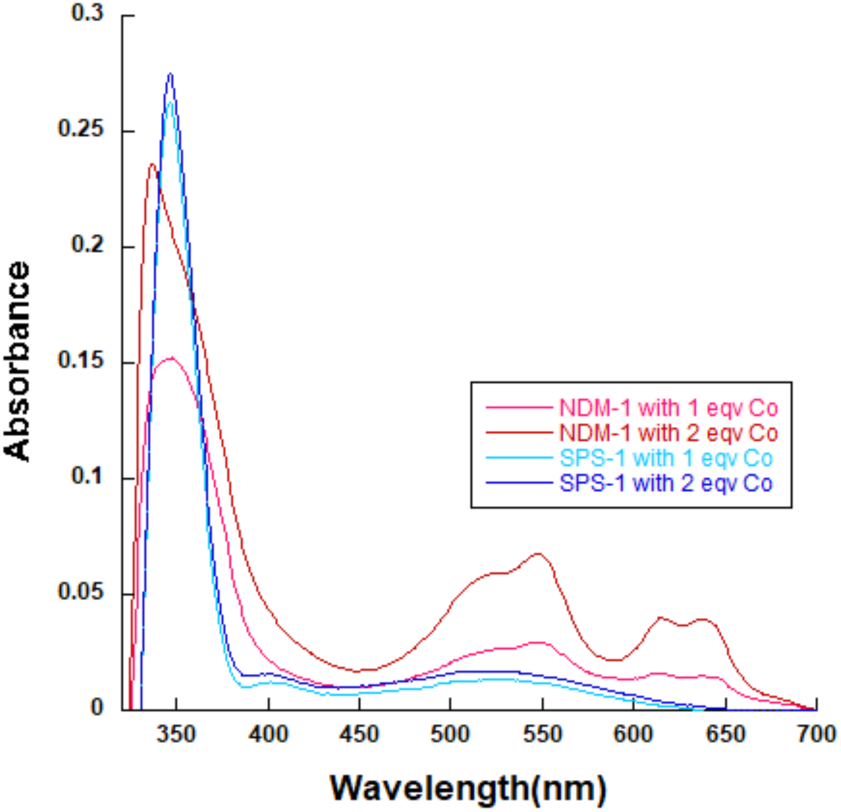
UV-vis spectra of apo-NDM and apo-SPS-1 treated with CoCl_2_. The apo-form protein (300 μM) with 50 mM HEPES, pH 6.8, containing 150 mM NaCl were used as blank in this experiment.

**NMR spectroscopy of Co (II)-substituted SPS-1.** To further probe the metal binding sites in SPS-1, ^1^H NMR spectra were obtained for Co (II)-substituted SPS-1 (Figure 4). As with the UV-Vis spectra above, Co (II)-substituted NDM-1 was used as a control. Previously for Co (II)-substituted NDM-1, we showed that resonances at 64, 72, and 79 ppm are due to solvent-exchangeable protons on His120, His122, and His189 in the Zn_1_ site, and resonances at 48, 107, and 165 ppm correspond to protons on Asp124, His250, and Cys208 in the Zn_2_ site, respectively (17). The spectrum of Co (II)-substituted SPS-1 revealed significant changes to the active site compared to NDM-1. Resonances at 249 and 217 ppm may be the geminal pair arising from the β-CH_2_ cysteine ligand coordinating the Zn_2_ site, although another weak signal was seen at 164 ppm that aligns well with the β-CH_2_ cysteine protons of Co (II)-substituted NDM-1. The SPS-1 data, therefore, suggest either two distinct conformations for Cys208, or that only two of the three resonances arise from Cys208 (25). Three relatively strong resonances were observed at 96, 74, and 54 ppm and can be assigned to the three Co (II)-bound histidine ligands (two from the Zn_1_ site and one from the Zn_2_ site). A spectrum of Co (II)-substituted SPS-1 containing 90% (v/v) D_2_O was collected to verify that these three resonances were solvent-exchangeable (Figure S2). The absence of these three resonances in the D_2_O spectrum further supports the hypothesis that they arise from the three Co (II)-coordinated histidine residues. Other resonances at 31 and 23 ppm do not appear to be solvent-exchangeable and may be due to either one or both α-CH_2_ aspartate protons. Of the remaining weak signals (66, 62, 48, and-27 ppm, see Fig. S2), most are likely contributions from second sphere interactions with the metal sites. In total, these results align well with the predicted two histidine ligands at the Zn_1_ site and one histidine at the Zn_2_ site and show distinct differences between the active site structures of SPS-1 and NDM-1.

**FIGURE 4.**
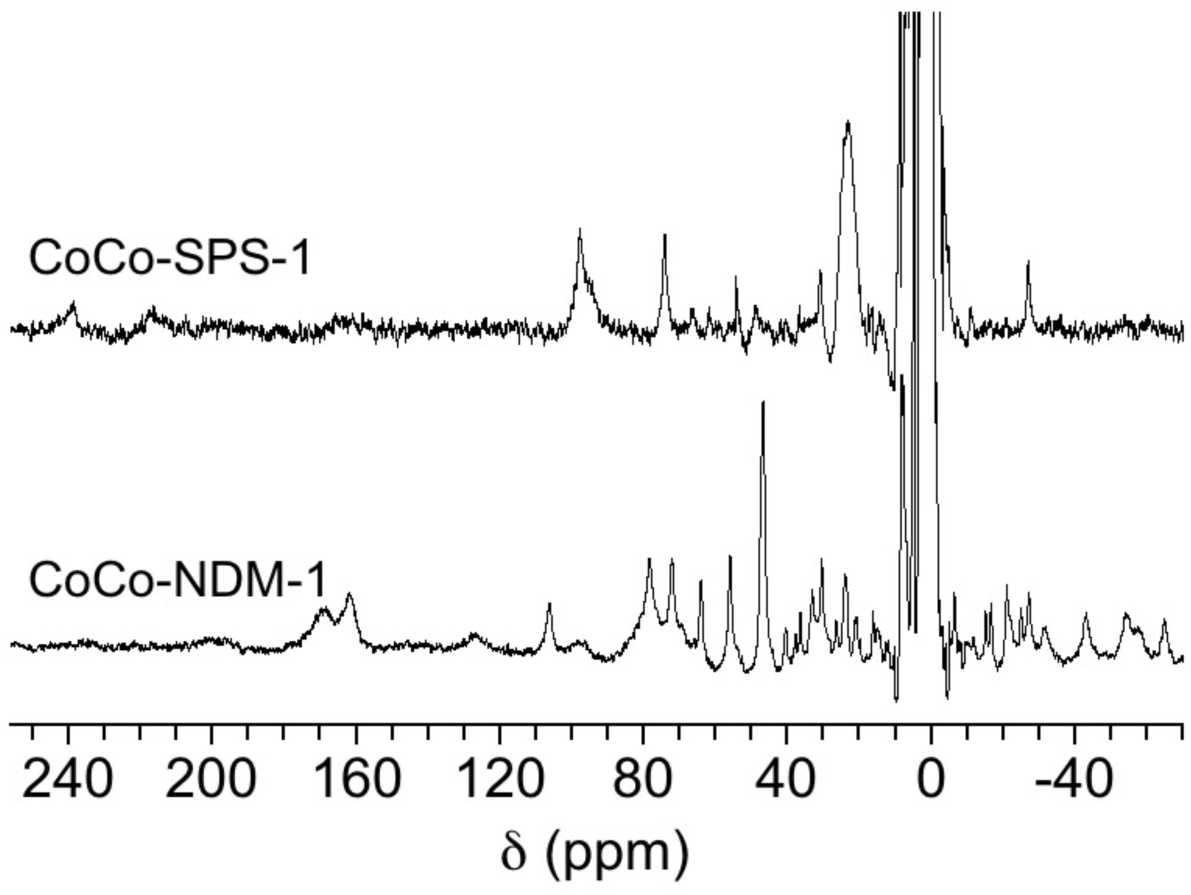
^1^H NMR spectra of Co (II)-substituted MβLs. 300 MHz ^1^H NMR spectra of Co (II)-substituted NDM-1 (1 mM) and SPS-1 (0.5 mM).

**EPR spectroscopy of metal-substituted SPS-1.** Continuous wave EPR spectra of CoCo-SPS-1 and CoCo-NDM-1, obtained in both parallel and perpendicular modes, are shown in Figure 5. Both perpendicular mode spectra are typical of high-spin Co (II)-substituted metalloproteins (26). Aside from modulation of line-shape, significant changes to the electronic environment of the Co (II) ions were not observed in these data. The sharp peak between 1600-1700 G in the diCo (II)-substituted NDM-1 spectrum was caused by a small fraction of Fe (III) contamination; a similar peak has been observed in previous EPR spectra of Co (II)-substituted NDM-1 (17). Co (II)-substituted NDM-1 also showed tightly-coupled, active site Co (II) ions in the parallel mode spectra (a deep negative feature near 800 G) (26). However, no such feature was observed in the spectrum of Co (II)-substituted SPS-1, indicating weaker coupling of active site Co (II) ions in SPS-1.

**FIGURE 5.**
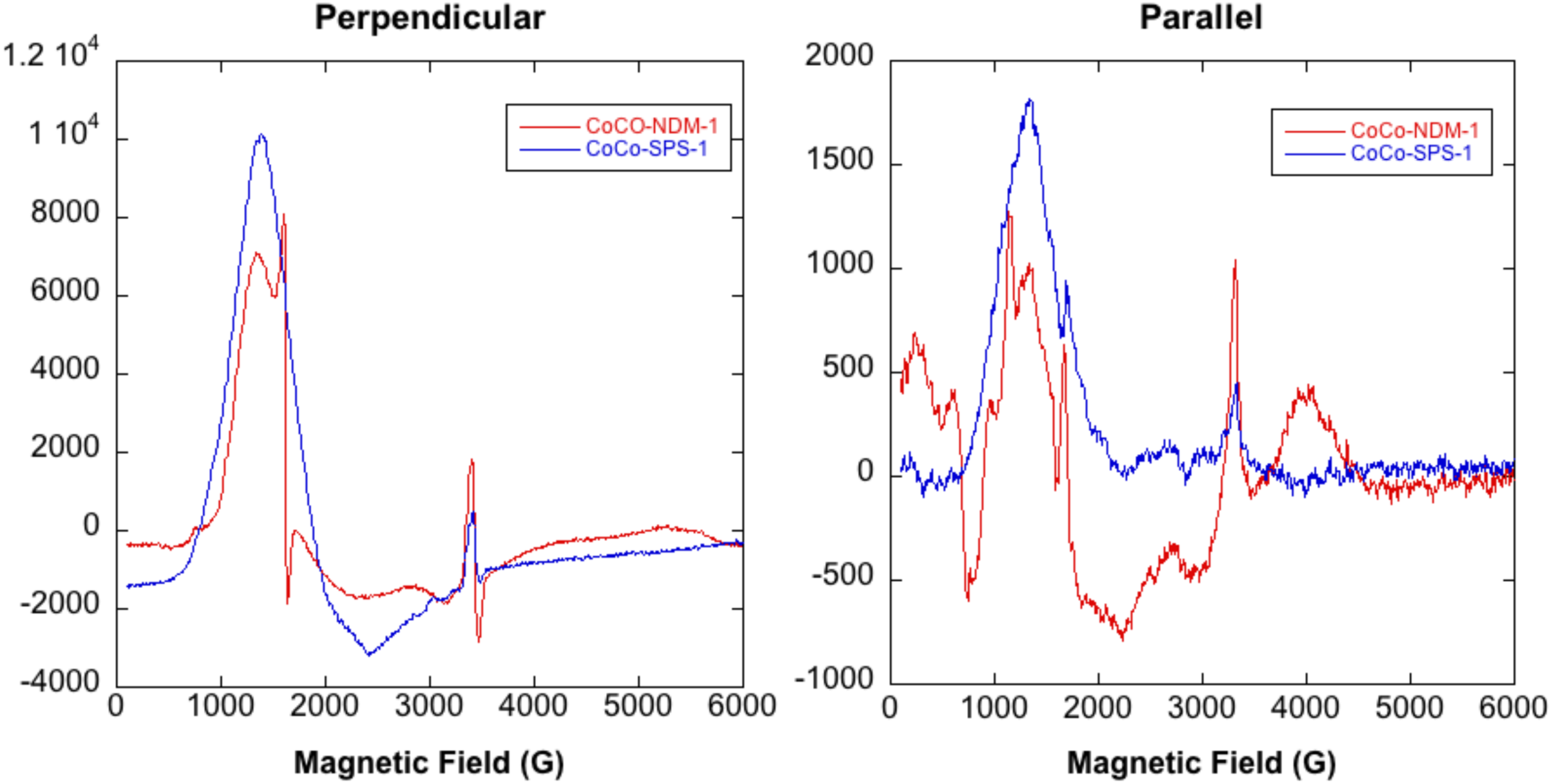
EPR spectra of Co (II)-substituted MβLs. Perpendicular and parallel mode, X-band EPR spectra of Co (II)-substituted NDM-1 SPS-1. The enzyme concentration in both samples was 0.9 mM.

**Crystal structure of SPS-1.** The structure of SPS-1 was solved to 1.70 Å resolution featuring an asymmetric unit containing a single protomer. Details of the crystallographic refinement are given in Table 2. Two Zn (II) ions are bound in the active site, and a third zinc ion is coordinated by the side chains of the His110 and Asp106 residues of the single protomer within the asymmetric unit and the side chains His110 and Asp106 of the-X,-Y,-Z+1/2 symmetry mate. This third Zn (II) ion assists in forming crystal contacts and is unlikely to be physiologically relevant. The SPS-1 structure (Figure 6 a,b) exhibits the canonical αββα metallo-β-lactamase fold with two separate β-sheets and interstrand loops facilitating coordination of the active site zinc ions. SPS-1 overlays with a 0.92 Å rmsd to SPM-1, indicating similar structures for both enzymes. The overlay of SPS-1 and NDM-1 is slightly poorer, exhibiting a 1.10 Å rmsd. Despite a slightly worse rmsd, SPS-1 exhibits high structural similarity to both SPM-1 and NDM-1. Similar to SPM-1, SPS-1 features a pair of extended α-helices (Figure 6 a,c), SPS-1 helices α3 and α4, located above the active site. In comparison, these helices are not found in NDM-1 (Figure 6d). In contrast, the β-hairpin loop found in NDM-1 is significantly truncated in both SPS-1 and SPM-1.

**Table 2:**
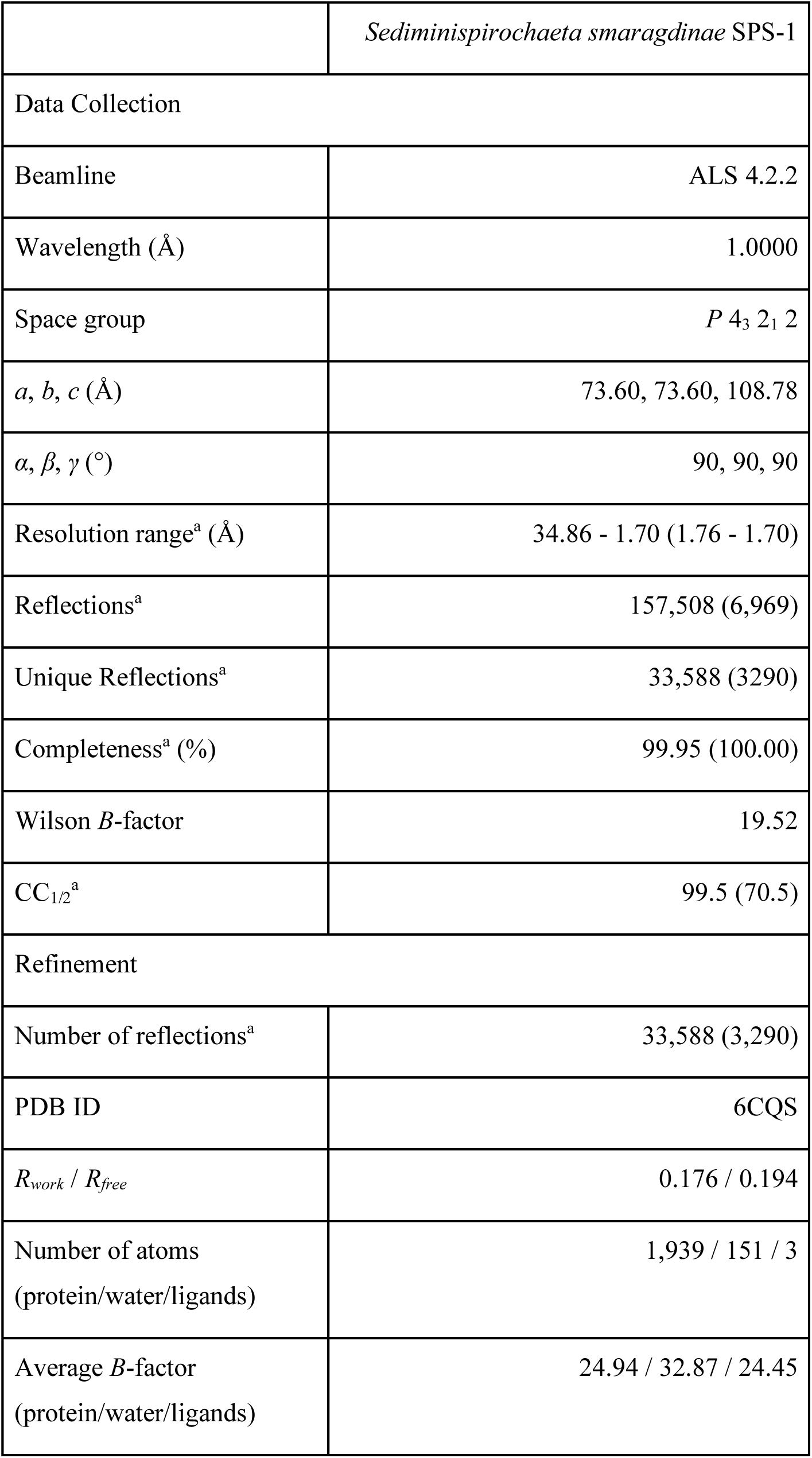
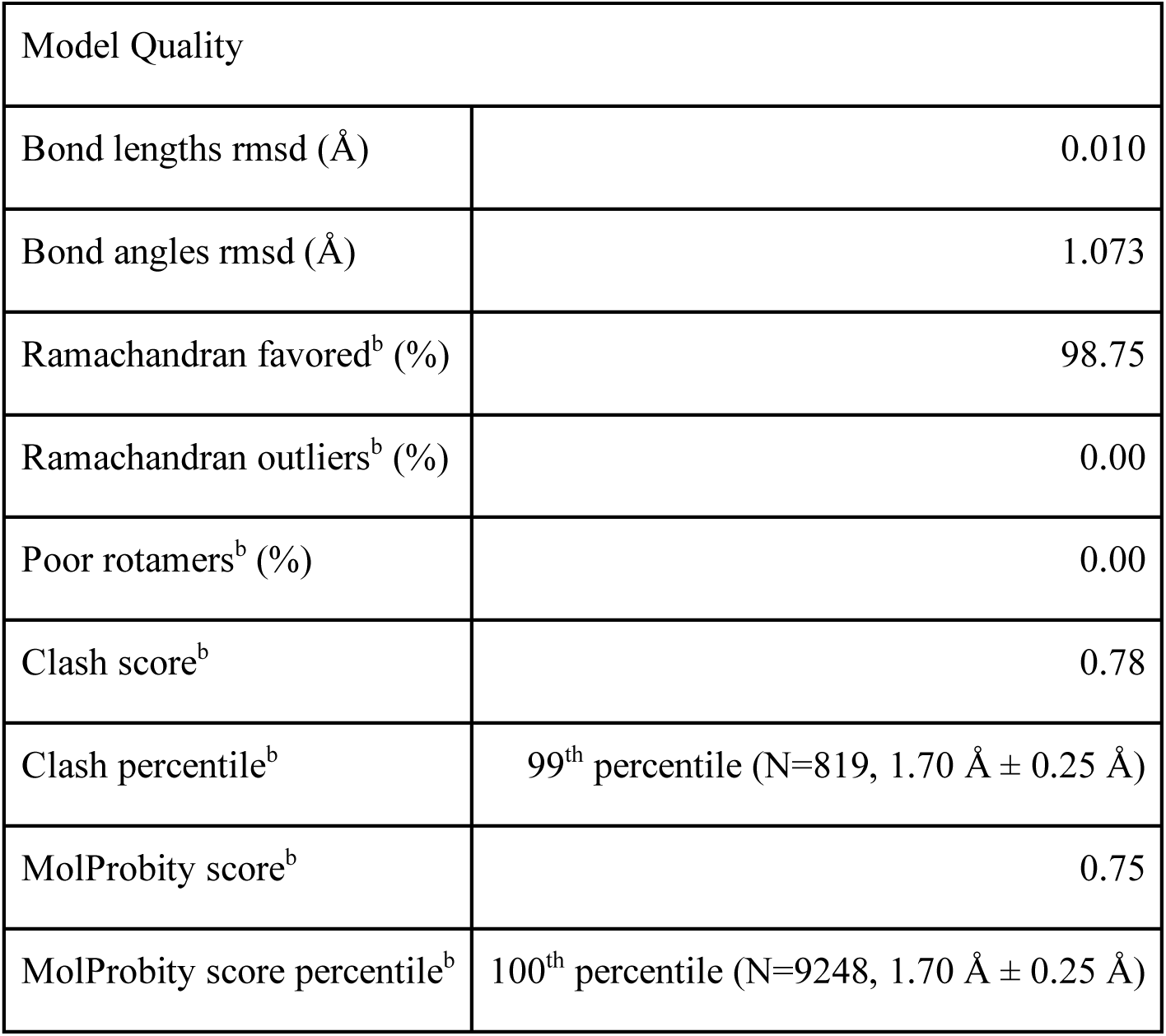
Crystallographic data on SPS-1

|  |  |
| --- | --- |
|  | *Sediminispirochaeta smaragdinae* SPS-1 |
| Data Collection |  |
| Beamline | ALS 4.2.2 |
| Wavelength (Å) | 1.0000 |
| Space group | $P 4_3 2_1 2$ |
| $a, b, c$ (Å) | 73.60, 73.60, 108.78 |
| $\alpha, \beta, \gamma$ (°) | 90, 90, 90 |
| Resolution range<sup>a</sup> (Å) | 34.86 - 1.70 (1.76 - 1.70) |
| Reflections<sup>a</sup> | 157,508 (6,969) |
| Unique Reflections<sup>a</sup> | 33,588 (3290) |
| Completeness<sup>a</sup> (%) | 99.95 (100.00) |
| Wilson $B$-factor | 19.52 |
| CC<sub>1/2</sub><sup>a</sup> | 99.5 (70.5) |
| Refinement |  |
| Number of reflections<sup>a</sup> | 33,588 (3,290) |
| PDB ID | 6CQS |
| $R_{work} / R_{free}$ | 0.176 / 0.194 |
| Number of atoms<br>(protein/water/ligands) | 1,939 / 151 / 3 |
| Average $B$-factor<br>(protein/water/ligands) | 24.94 / 32.87 / 24.45 |

| Model Quality |  |
| --- | --- |
| Bond lengths rmsd (Å) | 0.010 |
| Bond angles rmsd (Å) | 1.073 |
| Ramachandran favored<sup>b</sup> (%) | 98.75 |
| Ramachandran outliers<sup>b</sup> (%) | 0.00 |
| Poor rotamers<sup>b</sup> (%) | 0.00 |
| Clash score<sup>b</sup> | 0.78 |
| Clash percentile<sup>b</sup> | 99<sup>th</sup> percentile (N=819, 1.70 Å ± 0.25 Å) |
| MolProbity score<sup>b</sup> | 0.75 |
| MolProbity score percentile<sup>b</sup> | 100<sup>th</sup> percentile (N=9248, 1.70 Å ± 0.25 Å) |

<sup>a</sup>Values in parentheses represent data for the highest resolution shell.
<sup>b</sup>Calculated with MolProbity v4.4 (52)

**FIGURE 6.**
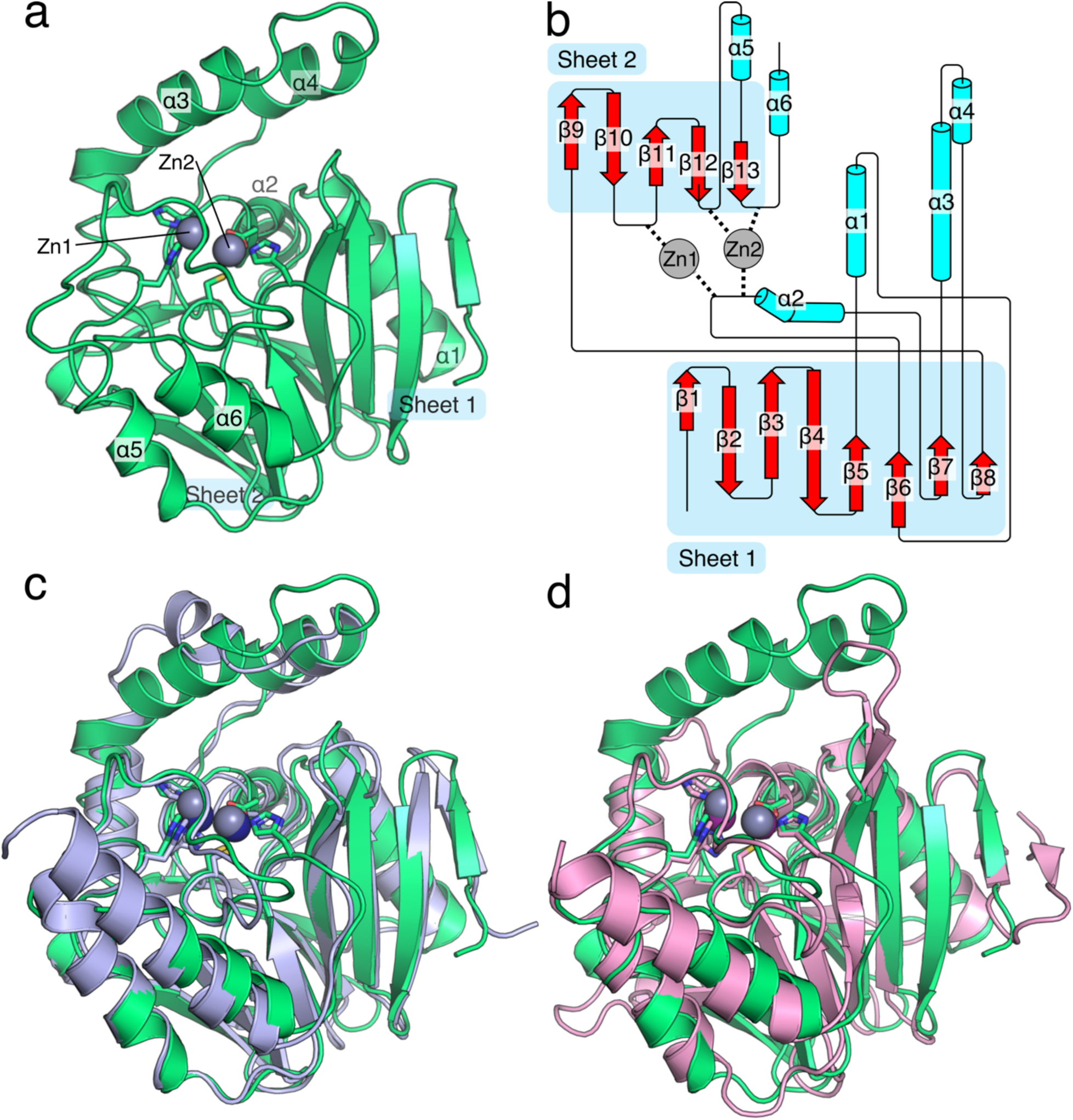
Overall structure of *S. smaragdinae* SPS-1. The structure of SPS-1 (a) features the canonical αββα metallo-β-lactamase fold with two zinc ions coordinated at the active site. Residues that coordinate zinc ions at the active site are shown as sticks, zinc ions are shown as spheres (*grey*). A cartoon schematic (b), generated in part by Pro-origami (54), indicates the organization of the protein and the contribution of loop residues to coordination of Zn_1_ and Zn_2_. An overlay of SPS-1 (*green*) with SPM-1 (*blue*, PDB ID 4BP0) highlights the similar overall structure (c). An overlay of SPS-1 (*green*) with NDM-1 (*pink*, PDB ID 4EXS) highlights a shortened SPS-1 β-hairpin loop between β-strands β3 and β4 and extended α-helices α3 and α4 above the active site for SPS-1 compared to NDM-1 (d). For (c) and (d) residues that coordinate zinc ions at the active site are shown as sticks, zinc ions are shown asspheres (SPS-1: *grey*, SPM-1: *navy blue*, NDM-1: *purple*).

**SPS-1 Active Site.** The active sites of SPS-1, SPM-1, and NDM-1 (Figure 7a) all feature two Zn (II) ions coordinated by the side chains of histidine, aspartate, or cysteine residues. The Zn_2_ site of SPS-1 (Figure 7b), SPM-1 (Figure 7c), and NDM-1 (Figure 7d) are nearly identical, each exhibiting coordination by an active site histidine (SPS-1 His245; SPM-1 His258; NDM-1 His250), aspartate (SPS-1 Asp100; SPM-1 Asp112; NDM-1 Asp124), and cysteine (SPS-1 Cys207; SPM-1 Cys216; NDM-1 Cys208) residues. While each protein features at least two histidine residues serving to coordinate Zn_1_ (SPS-1 His98 and His188; SPM-1 His110 and His197; NDM-1 His122 and His189), SPS-1 is missing the third coordinating histidine (SPM-1 His108; NDM-1 His120). In the place of His108/His120, SPS-1 harbors a glycine residue (Gly96). The substitution of a glycine at this position opens up substantial space in the SPS-1 active site below Zn_1_. As a result, SPS-1 features two water molecules that form a hydrogen bond network with SPS-1 active site residues. These two water molecules are featured clearly in the 2F_o_F_c_ map (Figure 8 a,b) and in an omit map generated by Polder (Figure 8 c,d) (27). While additional density in the omit map near Zn_1_ is likely the result of truncation artifacts in the Fourier series near the zinc ions, the omit map density clearly fits the proximal and distal water molecules (28). The water molecule proximal to Gly96 is held in place by a hydrogen bond to the backbone amide proton of Tyr97 and the active site water molecule proximal to Gly96. This proximal water molecule is further anchored by a hydrogen bond to the side chain NH_2_ of Asn192 and coordinates Zn_1_. The presence of the distal and proximal water molecules is made possible by a single amino acid substitution outside the primary coordination sphere. SPS-1 Thr187 is a bulky residue in comparison to SPM-1 Ala196 and NDM-1 Gly98, which results in a change in preferred rotamer for Asn192, thereby setting the stage for the water-mediated hydrogen bond network that situates the Gly96 distal water in position to coordinate Zn_1_.

**FIGURE 7.**
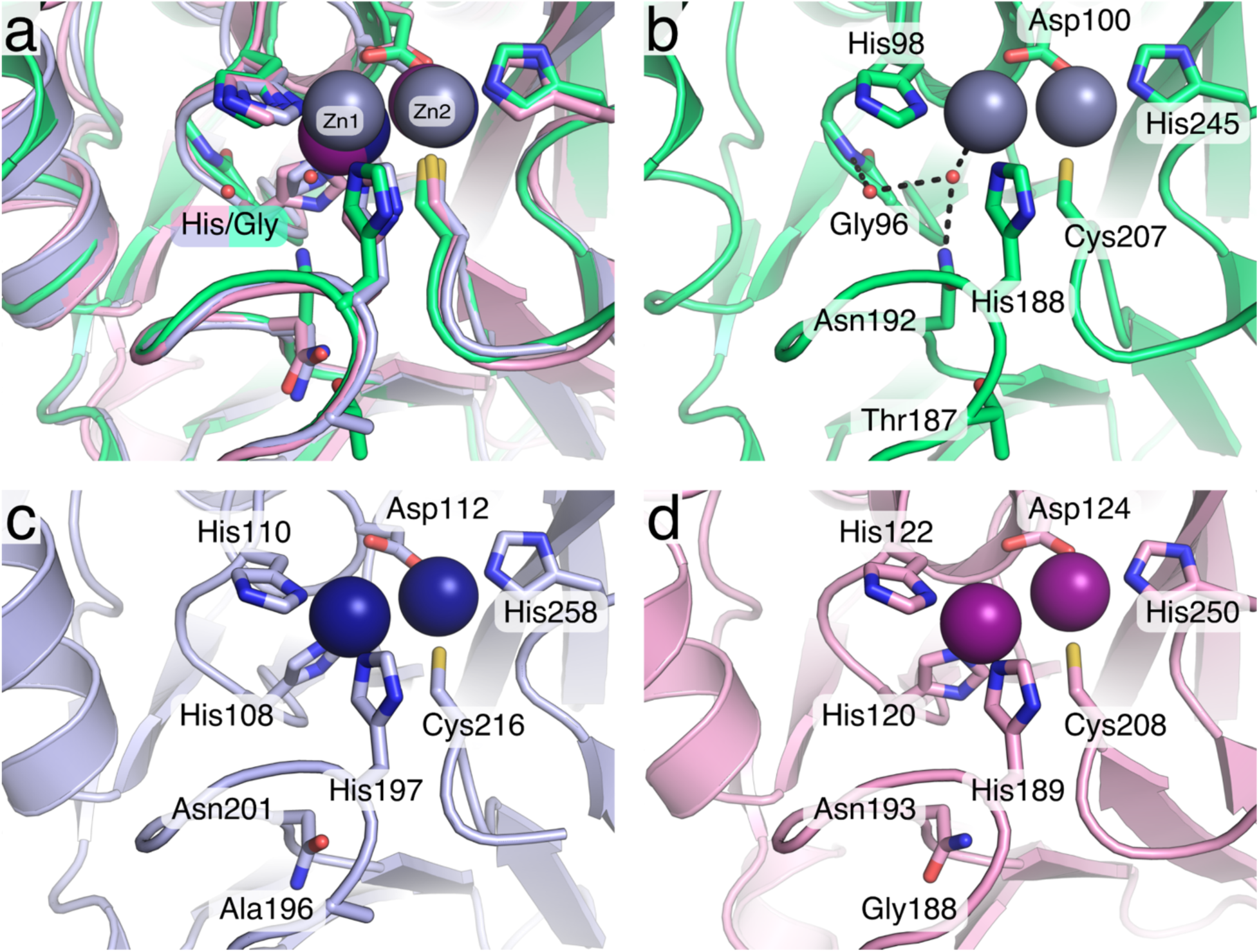
Organization of the *S. smaragdinae* SPS-1 active site. The active sites of SPS-1 (*green*), SPM-1 (*pink*, PDB ID 4BP0), and NDM-1 (*blue*, PDB ID 4EXS) are shown as an overlay (a). The active site of SPS-1 (b) identifies a water molecule that coordinates to Zn_1_ in place of a histidine from the canonical HxHxD motif and a second water molecule which completes the hydrogen bond network (*black dashed lines*) with Gly96 and Asn192. The active sites of SPM-1 (c) and NDM-1 (d) feature the full complement of expected ligands. Throughout, key residues are shown as sticks and identified by residue name and number, zinc ions are shown as spheres (SPS-1: *grey*, SPM-1: *navy blue*, NDM-1: *purple*).

**FIGURE 8.**
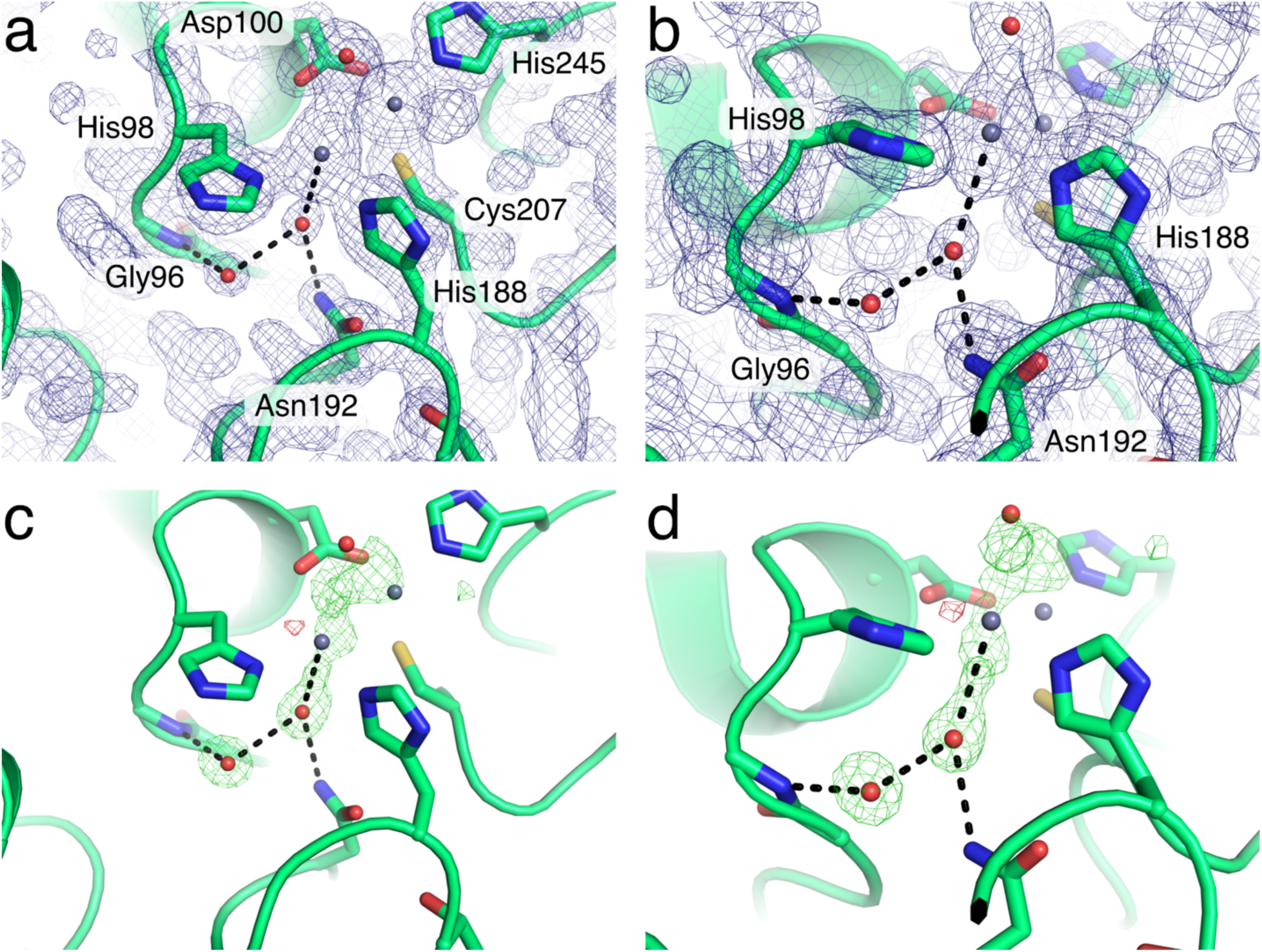
Electron density maps for the *S. smaragdinae* SPS-1 active site. Two views of the SPS-1 active site (a,b) with the 2F_o_-F_c_ map drawn at 1.5s. The same views of SPS-1 (c,d) are shown with an F_o_-F_c_ omit map generated by Polder (27) drawn at+/-4s with positive density colored green and negative density colored red. Throughout, key residues are shown as sticks, water molecules (*red*) and zinc ions (*grey*) are shown as small spheres (27).

The coordinating residues and the ions in the Zn_2_ sites harbor similar positions as the average distance between the Zn_2_ site zinc ions across the SPS-1, SPM-1, and NDM-1 structures are 0.57 ± 0.14 Å. In comparison, altered Zn_1_ coordination sphere has a significant effect on the location of Zn1 within the active site. The Zn_1_ site for SPS-1 diverges from those observed for SPM-1, NDM-1, and all other structurally characterized metallo-β-lactamases (1-3). While the position of Zn_2_ is relatively invariant, the position of SPS-1 Zn_1_ differs from SPM-1 Zn_1_ and NDM-1 Zn_1_ by 1.11 ± 0.01 Å (Figure S2). In comparison, the Zn_1_ sites of SPM-1 and NDM-1 differ by only 0.43 Å. The approximately 1 Å shift of SPS-1 Zn_1_ upward, away from Gly96 appears to be a direct consequence of the altered primary coordination sphere. Thus, differences in the primary (Gly96) and secondary (Thr187) coordination spheres elicit differences in the location of SPS-1 Zn_1_ in comparison to SPM-1 and NDM-1.

## DISCUSSION

A recent bioinformatics study revealed 279 unique, putative MβL genes, and initial biochemical and microbiological studies confirmed that 76 of the resulting proteins could be MβLs (6). One member of this group, SPS-1 from *S. smaragdinae*, was particularly interesting because it appeared to lack the consensus HXHXD motif found in all MβLs, except those belonging to the B2 subgroup (1-3). We were intrigued by the possibility of a novel metal binding site never before observed in the MβLs. Therefore, we endeavored to biochemically and structurally characterize SPS-1.

SPS-1 contains 276 amino acids and shares the highest sequence similarity with SPM-1 from *Pseudomonas aeruginosa* (Figure 1) (11). The overlay of the SPS-1 and SPM-1 structures reveal a 0.92 Å rmsd, indicating broad structural similarity (Figure 6 a,c). In addition, both SPS-1 and SPM-1 have a pair of helices located above the active site, and a similarly-positioned helix is not found in NDM-1 (Figure 6 a,c,d). Even though SPS-1 was sorted into the B1 phylogenetic class, it may also share some similarity with subgroup B2 MβLs, which contain one zinc in the active site (the consensus Zn_2_ site) and exhibit a narrow substrate spectrum (2). While NDM-1, VIM-2 and IMP-1 from B1 subclass can hydrolyze penems, cephems and carbapenems. CphA, Sfh-1, and ImiS from B2 subclass can only hydrolyze carbapenems (29). Surprisingly, SPS-1 exhibited significant activity against carbapenems, no activity against a penam, and weak-selective activity against cephem (Table 1). The most likely explanation for the absence of activity against ampicillin, Penicillin G, cefuroxime, and cephalexin is steric clashes with Trp48 and Thr146 located on the extended helices *α*3 and *α*4 in SPS-1 (Figures 6, 7). Overlaying SPS-1 onto structures of NDM-1 in complex with ampicillin (30), cephalexin (31), and cefuroxime (31) identifies that the elongated *α*3 and *α*4 helices of SPS-1 block the binding of substrates that contain large substituents in the 7’/8’ position in the *β*-lactam (Figure 9). In contrast, an overlay of SPS-1 onto structures of NDM-1 in complex with imipenem (32) and meropenem (16) (Figure 9), which lack the large substituent at the 7’ position, do not predict any steriv clashes between the penems and SPS-1. The 1 Å displacement of Zn (II) in the Zn_1_ site may also contribute to steric clash by forcing the substrate upward, but this movement would only exacerbate the clashes predicted for ampicillin, cephalexin, and cefuroxime.

**FIGURE 9.**
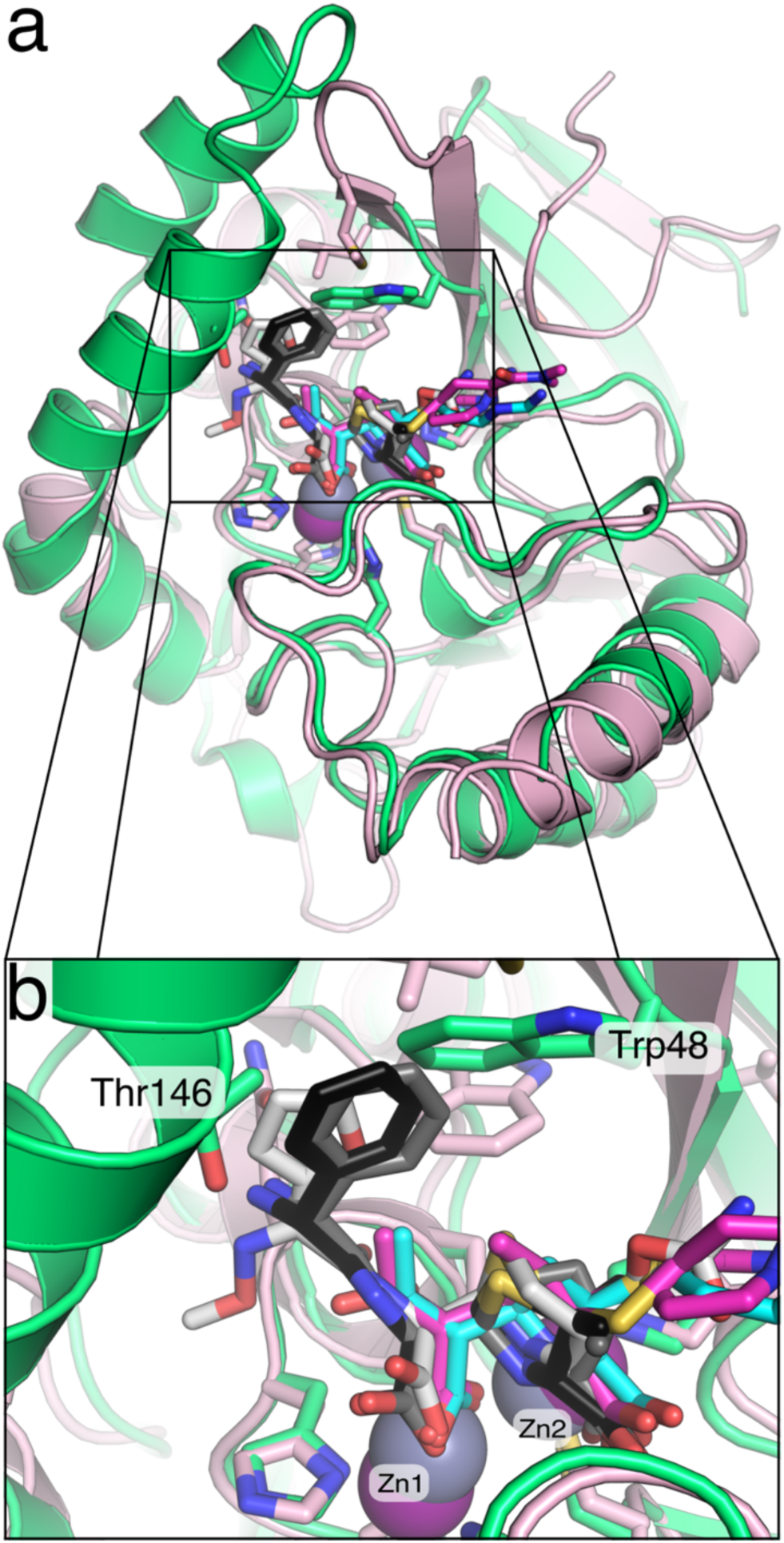
Overlay of *S. smaragdinae* SPS-1 and NDM-1 complexes with hydrolyzed antibiotics. The structure of SPS-1 (*green*, PDB ID 6CQS) was superimposed onto NDM-1 (*pink*, PDB ID 3Q6X) with SPS-1 Zn (II) ions (*grey*) and NDM-1 Zn (II) ions (*purple*) shown as spheres. Hydrolyzed antibiotics solved in complex with NDM-1 are shown as sticks including imipenem (*cyan*, PDB ID 5YPL), meropenem (*magenta*, 4EYL), ampicillin (*black*, PDB ID 3Q6X), cephalexin (g*rey*, PDB ID 4RL2), and cefuroxime (*white*, PDB ID 4RL0).

It is clear that the differences in relative position of Zn (II) at the Zn_1_ site does not in general lead to total loss of hydrolytic activity as SPS-1 is active against antibiotics, such as meropenem, which lack the phenyl or furyl substituents found in ampicillin, cephalexin, and cefuroxime (Figure 9). However, the different position and ligand environment of the Zn_1_ metal ion may explain the relatively larger *K*m values exhibited by SPS-1, as compared to those exhibited by NDM-1 and SPM-1 (Table 2). SPS-1 has two water molecules that form a hydrogen bond network, which accommodates the loss of a third coordinating histidine found in the Zn_1_ sites of all other B1 MBLs (Figure 7). For the dinuclear Zn (II)-containing MβLs, substrates are thought to bind with the *β*-lactam carbonyl oxygen interacting with Zn_1_ and the carboxylate group of the *β*-lactam interacting with Zn_2_ (1). Given the different active site structure in SPS-1, it is not surprising that the *K*m values for all tested substrates were higher, reflecting weaker substrate binding to a first approximation. This weaker binding is supported by ITC studies with competitive inhibitor captopril, which show that SPS-1 does not bind captopril. These results suggest that the different position of the Zn_1_ metal leads to the inability of captopril to bridge the Zn (II) ions in SPS-1. A surprising observation in the steady-state kinetic studies was that despite a different active site structure, higher *k*cat values were exhibited by SPS-1 towards carbapenems. Given the phylogenetic grouping of SPS-1 near the B2 MBLs, the positioning of the Zn_1_ metal above its position in other B1 MBLs, and the observed steady-state kinetic results, it is interesting to speculate that SPS-1 could be an intermediate enzyme between the dinuclear Zn (II)-containing MβLs (B1’s) and the mononuclear Zn (II)-containing MβLs (B2’s).

We have previously reported spectroscopic studies on several Co (II)-substituted MβLs, including NDM-1 (17), VIM-2 (4), IMP-1 (19), L1 (24), BcI (33), CcrA (18), Bla2 (34), and ImiS (8). In addition to better understanding the active sites of these enzymes, these studies have allowed us to probe metal binding to the MβLs. Like most reported MβLs, the Co (II)-substituted analog of SPS-1 can be prepared by a direct method in which native Zn (II) is removed with chelator and dialysis and the Co (II) analog is prepared by adding Co (II) directly to metal-free enzyme (4,17). The UV-Vis spectra of Co (II)-substituted SPS-1 reveal an intense peak at 346 nm, which can be attributed to a cysteine sulfur ligand to metal charge transfer band. The intensity of that peak is the greatest when SPS-1 has 1 equivalent of Co (II), demonstrating, as predicted from the crystal structure, that the first equivalent of metal binds to the Zn_2_ site. The longer wavelength portion of the SPS-1 spectra are different than those of most MβLs, which are often dominated by intense ligand field transitions between 450-600 nm from 4-coordinate Co (II) in the Zn_1_ site (see the control spectra of NDM-1 in Figure 3). Instead, the spectra of Co (II)-substituted SPS-1 show a broad transition in this part of the spectra, which can be also attributed to ligand field transitions. Similar spectra were reported for MβL L1. The intensity (*ε*_525nm_=56 M^-1^cm^-1^) of the ligand field transition (s) in Co (II)-substituted SPS-1 suggests that Co (II) in the Zn2 site is five-or six-coordinate. The small increase in this peak upon addition of a second equivalent of Co (II), which populates the Zn_1_ site, suggests that the coordination number of Co (II) in the Zn_1_ site is six.

^1^H NMR and EPR spectra of Co (II)-substituted SPS-1 nicely correlate with the crystal structure of the Zn (II)-containing enzyme. Three solvent-exchangeable peaks can be assigned to histidine residues bound to high spin Co (II), and other resonances, by comparison to spectra of Co (II)-MβLs, can be assigned to cysteine and aspartate residues. The broader peaks observed in the NMR spectra of Co (II)-substituted SPS-1, as compared to those in the spectra of Co (II)-substituted NDM-1, suggest a slower T_1e_ for the unpaired electrons on Co (II) and Co (II) ions that are at best weakly spin-coupled (35,36). The lack of spin coupling is clearly observed in the parallel mode EPR spectra of Co (II)-substituted SPS-1. The positioning of the metal ion in the Zn_1_ site is most likely the cause of the loss of coupling, and nicely explains the lack of binding by captopril.

An intriguing question regarding SPS-1 is why *S. smaragdinae*, specifically strain SEBR 4228^T^ from a water sample near a Congo offshore oilfield, produces an MβL. Spirochetes are spiral-shaped bacteria and are pathogenic, causing yaws, Lyme’s disease, relapsing fever, and syphilis (37). One common treatment for some spirochete infections is *β*-lactams (38), and it is possible that another spirochete strain transferred resistance to *S. smaragdinae* strain SEBR 4228^T^. SPS-1 from *S. smaragdinae* has not been identified in any patients and there are no reports of any human infections due to *S. smaragdinae*. According to Hatosy (39), there are four possible mechanisms to explain how bacteria in the ocean gain antibiotic resistance. The first prediction is that antibiotic-resistant bacteria migrate from land into marine environments. The second prediction is that antibiotics enter into the marine environment, causing bacteria to evolve antibiotic resistant phenotypes. However, both of these mechanisms seem unlikely due to dilution of the bacteria and antibiotics by the ocean. The third prediction is that antimicrobial compounds are being produced *in situ* by marine organisms, potentially to give these organisms a survival advantage. The fourth prediction is that the proteins conferring antibiotic resistance phenotypes mutated from a protein that has a different function *in vivo*. SPS-1, like other MβLs, adopts a characteristic αββα fold. This αββα fold is found in a larger superfamily of metalloenzymes, such as glyoxalase II, arylsulfatase, and RNA processing enzymes (40) (41), with diverse biological functions. It is important to note that the evolution of one of these enzymes into a MβL would likely require the presence of *β*-lactams to drive evolution.

Despite the question of why this spirochete produces a MβL, SPS-1 is a unique MβL with a novel metal binding site. SPS-1 exhibits substrate specificity similar to the B2 MβLs, yet SPS-1 can hydrolyze *β*-lactams other than carbapenems. The novel ligand set for the SPS-1 active site provides a glimpse at possible mutations in B1 MβLs that can still allow for β-lactam hydrolysis, suggesting that broader diversity is possible for this family of enzymes. The uncoupling of the Zn_1_ and Zn_2_ metal sites identifies SPS-1 as a possible intermediate between the dinuclear Zn (II)-containing B1 MβLs and mononuclear Zn (II)-containing B2-MβLs. Additional studies are needed to better understand how the different metal center in SPS-1 might affect the mechanism of *β*-lactam hydrolysis and the interaction with current and future inhibitor scaffolds.

## EXPERIMENTAL PROCEDURES

**Phylogenetic analysis of SPS-1.** The protein sequences of selective class B β-lactamases (MβLs), as well as SPS-1, were downloaded from NCBI Genbank and analyzed using Phylogeny.fr (http://www.phylogeny.fr/) (10,42).

**Cloning, over-expression, and purification of SPS-1.** A codon-optimized SPS-1 sequence (residues 32-276) fused with a N-terminal Tobacco Etch Virus (TEV) cleavage site was synthesized by Genscript Biotech Corporation. The synthesized fragment was sub-cloned into pET28a between the restriction sites *Nde*I and *Xho*I. The resulting over-expression plasmid, pET28a-SPS-1, was transformed into *Escherichi. coli* EXPRESS BL21 (DE3) Chemically Competent cells (Lucigen). A single colony was transferred into 50 mL of lysogeny broth containing 50 μg-mL kanamycin, and the culture was shaken overnight at 37 °C. The overnight culture (10 mL) was used to inoculate four 1 L LB cultures containing 50 μg-mL kanamycin. The resulting culture was grown at 37 °C with a shaking speed of 220 rpm until an OD_600_ of 0.6 was reached. Protein production was induced by addition of IPTG to 500 μM. At time of induction, cultures were supplemented with Zn (II) with addition of ZnCl_2_ to 100 μM. Cultures were shaken overnight at 18 °C and then harvested by centrifugation for 10 min at 8,000 xg. Cell pellets were resuspended in 50 mL of 50 mM HEPES, pH 7.5, containing 500 mM NaCl. Cells were lysed by passing the mixture two times through a French Press at a pressure between 15,000-20,000 psi. The insoluble components were removed by centrifugation for 1 hour at 32,000 xg. The supernatant was loaded onto HisTrap HP (5 mL, GE) column and washed with 10 volumes of wash buffer containing 50 mM HEPES, pH 7.5, 500 mM NaCl, and 50 mM imidazole. Bound proteins were eluted using elution buffer (wash buffer with 500 mM imidazole). TEV protease was added at a protease to target protein ratio of 1:100 (w/w). The eluted proteins were dialyzed against 1L of 50 mM HEPES, pH 7.5, containing 500 mM NaCl. After overnight dialysis, the truncated SPS-1 protein was re-purified by passage through a HisTrap column, which removed the His_6_-TEV and His_6_-tag.

Purified recombinant SPS-1 (before and after TEV digestion) was analyzed by sodium dodecyl sulfate-polyacrylamide gel electrophoresis (SDS-PAGE) using Fisher EZ-Gel solution (12.5%), a Mini-PROTEAN System (Bio-Rad Laboratories), and Coomassie Blue stain. Protein concentrations were determined by UV-Vis absorbance at 280 nm using the ε_280nm_=35,535 M^-1^ cm^-1^ for SPS-1 protein containing the His_6_-tag and ε280nm=34,045 M^-1^ cm^-1^ for the SPS-1 protein without the His_6_-tag. These extinction coefficients were calculated using ExPASY ProtParam (13).

**MALDI-TOF mass spectrometry.** After dialysis against 10 mM ammonium acetate, pH 7.5, 30 μM tag-free SPS-1 was mixed with sinapinic acid matrix at a ratio of 1:1. The resulting sample was analyzed using a Bruker AutoFlex MALDI-TOF mass spectrometer (43).

**Fluorescence spectroscopy.** Fluorescence emission spectra of SPS-1 were acquired on a Perkin Elmer Luminescence spectrometer (Model LS-55) using 2 μM SPS-1 in 50 mM HEPES, pH 7.5). Spectra were acquired using an excitation wavelength of 280 nm and an emission spectrum window from 300 to 500 nm.

**Metal Analyses.** The zinc content of purified SPS-1 samples were determined using a Perkin-Elmer Optima 7300V inductively coupled plasma spectrometer with optical emission spectroscopy (ICP-OES) detection. Protein samples, used without modification after purification, were diluted to 5 μM with 10 mM ammonium acetate, pH 7.5. Calibration curves were generated using serial dilutions of FisherBrand Zn metal standard ranging from 0 to 16 μM, and the emission line at 202.548 nm was chosen for zinc.

**Steady-state kinetics.** Steady-state kinetic studies were performed using a Synergy HT Multi-Mode Microplate Reader (Biotek) at 37°C in 50 mM HEPES, pH 7.5, containing 150 mM NaCl and 10 μM ZnCI_2_. The changes in molar absorptivity (Δε, M^-1^cm^-1^) used to quantitate products were: chromacef, Δε_442nm_=18,600; cephalothin, Δε_265nm_ =-8,790; imipenem, Δε_300_ =-9,000; meropenem, Δε_293_ =-7,600; ampicillin, Δε_235_ =-809; and penicillin G, Δε_235_ =-936; cefuroxime, Δε_260_ =-5,860; and cephalexin, Δε_260_ =-7,750. (11) Initial rates (within 18 seconds) versus substrate concentrations were fitted to the Michaelis-Menten equation to determine the steady-kinetic constants *K*_m_ and *k*_cat_ using Prism 7 (Graphpad software).

**Isothermal titration calorimetry.** Isothermal titration calorimetry (ITC) experiments were carried out using a Nano ITC System (TA Instruments-Waters LLC., USA) with a 500 μL cell. All experiments were performed at 25 °C. SPS-1 and NDM-1 were prepared by diluting the stock solutions with 20 mM HEPES, pH 7.5, containing 150 mM NaCl. *L*-Captopril (Acros Organics) was dissolved in the same buffer. The ITC cell was filled with 50 μM of NDM-1 or SPS-1 (300 μL), and the enzyme solutions were titrated with 250 μM captopril. The injection volume was 50 μL, and the time between two injections was 210 s. K_d_ values were determined by using NanoAnalyze (TA Instruments-Waters LLC., USA).

**Preparation of metal-free proteins.** Metal-free forms of SPS-1 and NDM-1 were prepared through metal chelation. Each purified protein (diluted to 50 μM in 20 mL) was placed in a 10 kDa molecular weight cutoff dialysis bag. These samples were dialyzed versus dialysis buffers for 4 hours at 4 °C before changing to the subsequent dialysis buffer to include a series of eleven dialysis steps: First, two changes of 50 mM HEPES, pH 6.8, containing 150 mM NaCl with 2 mM EDTA, then three changes of 50 mM HEPES, pH 6.8, containing 150 mM NaCl. The resulting metal-free enzymes were analyzed by using SDS-PAGE and UV-vis and concentrated by using an Amicon Ultra 15 mL centrifugal filter (10K).

**UV-Vis spectroscopy.** Metal-free SPS-1 and NDM-1 were diluted to 300 μM with 50 mM HEPES, pH 6.8, containing 150 mM NaCl, and 1 mM TCEP. One or two molar equivalents of CoCl_2_ were added to the protein samples from a 50 mM stock solution. The resulting mono-Co (II)-substituted or di-Co (II)-substituted protein samples were placed in 500 μL quartz cuvettes, and UV-Vis spectra were collected on a PerkinElmer Lambda 750 UV-VIS-NIR Spectrometer that measured absorbance between 300 to 700 nm at 25 °C. The blank spectrum of 300 μM apo-SPS-1 or apo-NDM-1 in 50 mM HEPES, pH 6.8, containing 150 mM NaCl and 1 mM TCEP was used to generate difference spectra. All data were normalized to the absorbance at 700 nm.

**NMR spectroscopy.** Metal-free SPS-1 was concentrated to ∽0.5 mM with an AmiconUltra-15 Centrifugal unit and an Ultracel-10 membrane. To the concentrated protein, 2 equivalents of CoCl_2_ were added in the buffer containing 50 mM HEPES, pH 6.8, containing 150 mM NaCl and 10% (v-v) D_2_O. Spectra were collected at 292 K on a Bruker ASX300 (BBI probe) NMR spectrometer operating at a frequency of 300.16 MHz. Spectra were collected using a frequency switching method, applying a long low power (270 ms) pulse centered at the water frequency, followed by a high power 3 μs pulse centered at either 200 or 90 ppm to ensure all resonances were detected (44). This method allowed for suppression of the water signal with enhancement of severely hyperfine-shifted resonances. Spectra consisted of 120,000 transients of 16k data points over a 333 ppm spectral window (*τ*_aq_ ∽51 ms). Signal averaging took ∽12 hours per spectrum.

**EPR spectroscopy.** EPR samples containing SPS-1 or NDM-1 included approximately 10% (v-v) glycerol as glassing agent. Samples were loaded into EPR tubes and degassed by repeated evacuation-purgation with N_2_ prior to data collection. Spectra were collected on a Bruker EMX EPR spectrometer, equipped with an ER4116-DM dual mode resonator (9.37 GHz, parallel; 9.62 GHz, perpendicular). The data in EPR figures were scaled so that the X-axes matched (perpendicular mode field values were scaled by 9.37/9.62). Temperature control was accomplished using an Oxford ESR900 cryostat and temperature controller (4.5 K). Other spectral conditions included: microwave power=0.2 mW; field modulation=10 G (100 kHz); receiver gain=10^4^; time constant/conversion time=41 ms.

***Crystallization, diffraction data collection, and structure refinement.*** SPS-1 was crystallized using 96-well sitting drop vapor diffusion screens prepared using a Phoenix crystallization robot (Art Robbins, USA). The sitting drops contained 400 nL of 1.45 mM SPS-1 and 400 nL of crystallization condition. Six crystallization screens were used: MCSG 1-4 by Microlytic and JCSG and PACT++ by JBScreen. Diffraction quality crystals formed within 2 weeks in a condition consisting of 0.2 M magnesium chloride, 0.1 M sodium cacodylate, pH 6.5, and 50% (v/v) PEG 200. For crystal harvesting, MiTeGen MicroLoops were used to harvest crystals that had been cryoprotected with LV CryoOil and then immediately frozen in liquid nitrogen. Crystals were sent to the Advanced Light Source (ALS) at Lawrence Berkeley National Laboratory where X-ray diffraction data were collected. Data were integrated and scaled using XDS (45) and SCALA (46). Data resolution cut-offs were determined using a combination of criteria including completeness and CC_1/2_. Molecular replacement was conducted with PHASER (47) using a poly-alanine search model produced from the structure of SPM-1 (PDB ID 4BP0) (48). The SPS-1 amino acid sequence was then docked to the molecular replacement solution. Iterative model building and refinement were conducted using COOT (49) and PHENIX (50) using isotropic individual *B*-factors and TLS refinement (51). Stereochemical and geometric quality analyses were performed with MolProbity version 4.4 (52). The final model and structure factors were deposited in the Protein Data Bank under PDB ID 6CQS.

## Acknowledgements

The authors acknowledging funding from the National Institutes of Health (R01 GM111926 to DLT, RAB, RCP, and MWC. RCP acknowledges support from Miami University through the Robert H. and Nancy J. Blayney Professorship. The Advanced Light Source is supported by the US Department of Energy under contract number DE-AC03-76SF00098 at Lawrence Berkeley National Laboratory. This work was also supported in part by the National Institutes of Health, National Institute of General Medical Sciences, and National Institute of Allergy and Infectious Disease under award numbers R01AI100560 (to RAB), R01AI063517 (to RAB), and R01AI072219 (to RAB). This study was also supported in part by funds and/or facilities provided by the Cleveland Department of Veterans Affairs, Award Number 1I01BX001974 (to RAB) from the Biomedical Laboratory Research & Development Service of the VA Office of Research and Development and the Geriatric Research Education and Clinical Center VISN 10 (to RAB). The content is solely the responsibility of the authors and does not necessarily represent the official views of the National Institutes of Health or the Department of Veterans Affairs.

## Author Contributions

Cloning, protein expression and purification, ITC, and fluorescence experitments were conducted by ZC, CM, KM, EC, HZ, RK, and SF. X-ray crystallography data collection and structure solution were performed by JVP, JCN, and RCP. MWC and RCP drafted the manuscript with input from the co-authors. All authors approved the final version of the manuscript.

